# Exploring *Alu*-Driven DNA Transductions in the Primate Genomes

**DOI:** 10.1101/2024.04.29.591526

**Authors:** Reza Halabian, Jessica M. Storer, Savannah J. Hoyt, Gabrielle A. Hartley, Jürgen Brosius, Rachel J. O’Neill, Wojciech Makałowski

## Abstract

Long terminal repeats (LTRs) and non-LTRs retrotransposons, aka retroelements, collectively occupy a substantial part of the human genome. Certain non-LTR retroelements, such as L1 and SVA, have the potential for DNA transduction, which involves the concurrent mobilization of flanking non-transposon DNA during retrotransposition. These events can be detected by computational approaches. Despite being the most abundant short interspersed sequences (SINEs) that are still active within the genomes of humans and other primates, the transduction rate caused by *Alu* sequences remains unexplored. Therefore, we conducted an analysis to address this research gap and utilized an in-house program to probe for the presence of *Alu*-related transductions in the human genome. We analyzed 118,489 full-length *Alu*Y subfamilies annotated within the first complete human reference genome, T2T-CHM13. For comparative insights, we extended our exploration to two non-human primate genomes, the chimpanzee and the rhesus monkey. After manual curation, our findings did not confirm any *Alu*-mediated transductions, whose source genes are, unlike L1 or SVA, transcribed by RNA polymerase III, implying that they are infrequent or possibly absent not only in the human but also in chimpanzee and rhesus monkey genomes. Although we identified loci in which the 3’ Target Site Duplication (TSD) was located distantly from the retrotransposed *Alu*Ys, a transduction hallmark, our study could not find further support for such events. The observation of these instances can be explained by the incorporation of other nucleotides into the poly(A) tails in conjunction with polymerase slippage.

## 1. Introduction

Discovered by Barbara McClintock in the 1940s, mobile elements [1], often termed transposable or transposed elements (TEs), are present in most, if not all eukaryotic genomes [2]. In the human genome, the discernible TEs contribute to about 46% or over 1.4 Gb of the sequence [3]. Based on the transposition mode, TEs have been divided into Type I elements that are mobilized via a “copy-and-paste” mechanism and Type II elements that move via a “cut-and-paste” manner [4]. Notably, not all copy-and-paste TEs can give rise to new copies, instead, there are only a limited number of master, source, or founder elements that spread their copies within genomes [5]. Therefore, most TEs are not transposable but instead only transposed elements [6]. Whether a TE is transposable presumably depends on retention of intact internal open reading frames and expression, particularly in the germ line. While for the L1 elements, *bona fide* source genes had been identified [7, 8], there is only sparse information on *Alu* source genes [9-12]. Autonomous elements, such as L1s, encode most of the gene products that are required for their activity, while non-autonomous elements, such as SVAs and *Alu*s rely on the machinery of autonomous elements to proliferate within genomes [13-15]. TE-families are subject to regular biological processes, including birth and death, i.e., a specific family or a subfamily of a TE appears in the genome and goes extinct after a period of activity. Consequently, only a small number of TE families are active within a certain evolutionary period. For instance, currently, only three non-LTR retroelements seem to be active in humans. These include one autonomous family (L1) and two non-autonomous families (*Alu* and SVA) [16] while older elements, such as mammalian-wide repeats (MIR), are still present and discernible, but their source genes are inactive [17, 18]. Interestingly, all these TEs belong to type I and consequently, *Alu*s and SVAs most likely employ the L1 molecular machinery for their retrotransposition. Although there are numerous sequences related to Type II transposons in the human genome, there is no evidence of their recent activity in primates [19].

At least by their abundance, TEs leave a significant legacy behind by influencing the evolution of genomes, including genome structure, gene evolution, and gene expression [20]. Jumping of TEs seems to be random, although there are some preferences concerning the genomic context into which the individual elements insert [21-24]. Nevertheless, in most cases, insertion of a TE into a new location has either a neutral or negative effect on the host genome in agreement with the neutral theory of evolution [25]. In fact, initial observations of the impact of retroposons on the human genome were made because of the disease phenotypes they caused [26, 27]. Early examples where a TE could transduce a piece of DNA unrelated to the TE’s original (source) locus to the new integration site (Figure 1) can be found in the literature [28-31]. In 1999 John Moran and colleagues demonstrated that active L1 elements are capable of DNA transductions *in vitro* [32]. Following these observations, several large-scale computational studies indicated the extent of the phenomenon at the genomic level [33-35]. While the potential evolutionary consequences of this process were noted [36], none of the studies confirmed exon shuffling caused by DNA transductions at fixed loci in the human genome. Unexpectedly, such a confirmation came from the studies of another active primate transposon, namely the SVA element [37].

**Figure 1.**
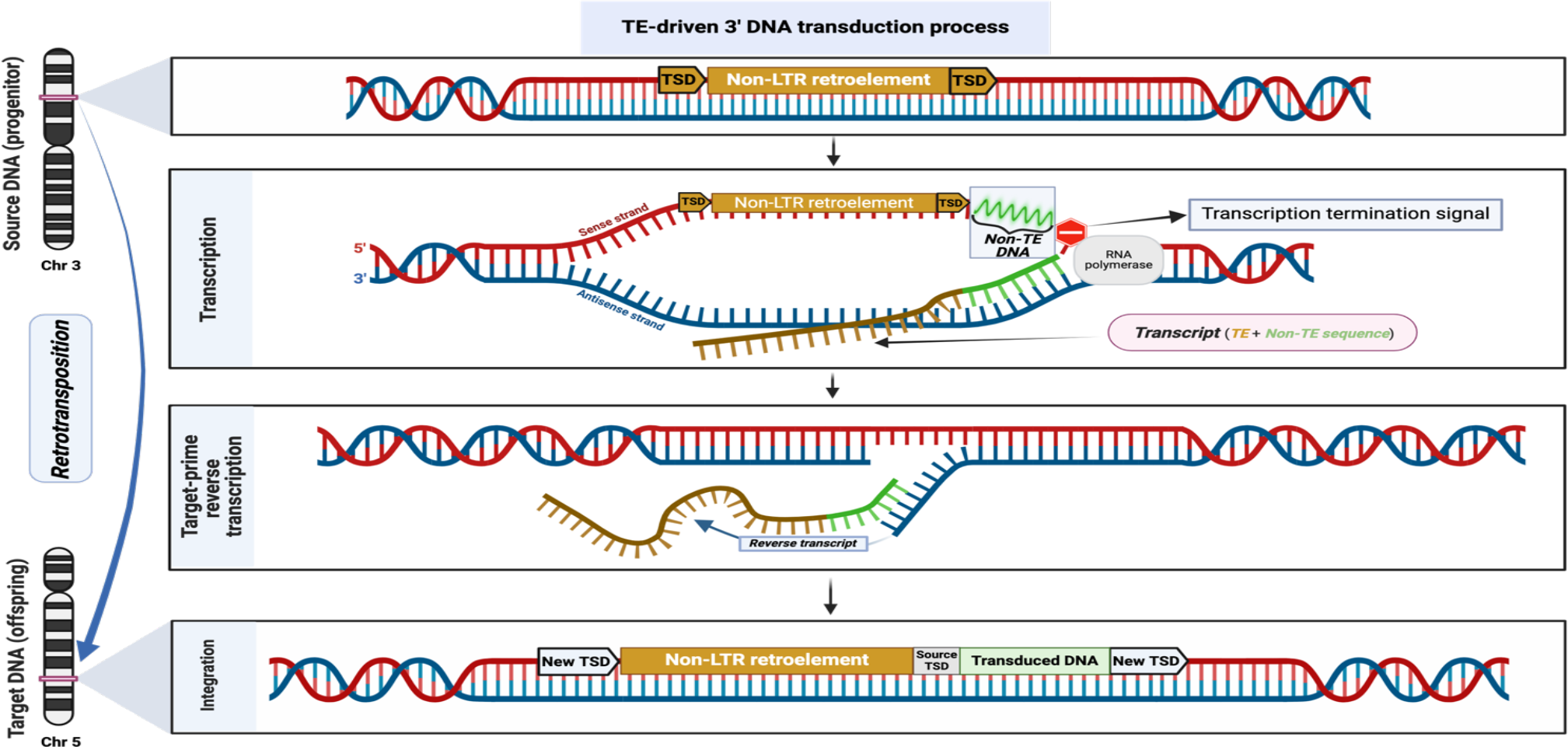
TE-driven DNA transduction. The cartoon illustrates a transduction process induced by a non-LTR retroelement. In the case of *Alu*s, the transcription termination motif is a poly T sequence with a minimum length of four nucleotides [38]. Hypothetically, if this polyT is located far from the source *Alu*, the resulting transcript comprises the original *Alu* and non-*Alu* sequence. This chimeric transcript has the potential to be inserted in a new genomic locus, giving rise to transduction.

Thus far, there are no systematic studies of *Alu*-driven DNA transductions. However, Kojima reported recent *Alu* monomer activity and one of these events was accompanied by a short DNA transduction [39]. Recently, Hoyt et al. reported that *Alu*-mediated DNA transduction occurred in the origin of AluSx-WaluSat, one of the recent composite elements in the human genome [3]. Consequently, we were interested in finding out the scale of *Alu*-driven DNA transductions in primates with a special focus on the human genome. Interestingly, despite over a million copies of *Alu* elements in the human genome, we could not confirm any evidence supporting the involvement of *Alu*s in the transduction process. However, it must be stressed that our investigation concentrated on *Alu*Ys, and we have applied very strict criteria throughout the analysis.

## 2. Materials and methods

### 2.1 Genome assemblies and annotations

We obtained the human reference genome (chm13v2.0.fa, accessed on August 15^th^, 2022) from the GitHub page of the T2T consortium (https://github.com/marbl/CHM13). Moreover, the RepeatMasker (CHM13v2.0_RM-2022MAR23.out) and segmental duplication (T2T-CHM13v2.SDs.bed) annotations were downloaded from the same repository on the same date. The genome of the chimpanzee, panTro6, (https://hgdownload.soe.ucsc.edu/goldenPath/panTro6/bigZips/panTro6.fa.gz) and the RepeatMasker annotation (https://hgdownload.soe.ucsc.edu/goldenPath/panTro6/bigZips/panTro6.fa.out.gz) were obtained on 12.12.2022. Finally, we accessed two files related to the rhesus monkey (rheMac10) on 22.12.2022 from the following repositories: the reference genome (https://hgdownload.soe.ucsc.edu/goldenPath/rheMac10/bigZips/rheMac10.fa.gz) and the RepeatMasker annotation (https://hgdownload.soe.ucsc.edu/goldenPath/rheMac10/bigZips/rheMac10.fa.out.gz).

### 2.2 Transduction identification and validation

Due to their different mode of transcription (RNA polymerase III), the existing tools for profiling 3’ transductions mediated by L1s seem inefficient for *Alu*s, as they often yield many false positives. Specifically, these tools lack a built-in verification step, requiring users to validate transductions by employing additional strategies. Hence, we developed a computational pipeline using a set of custom-built Python scripts to detect and confirm *Alu* transduction events more accurately (Figure 2). This method considers the transcriptional termination signal for pol III (a stretch of at least four thymine residues) during the validation step [38]. In addition, the method identifies the sequence and coordinates of target site duplications (TSDs) more precisely by constructing k-mers from segments around each *Alu* sequence.

**Figure 2.**
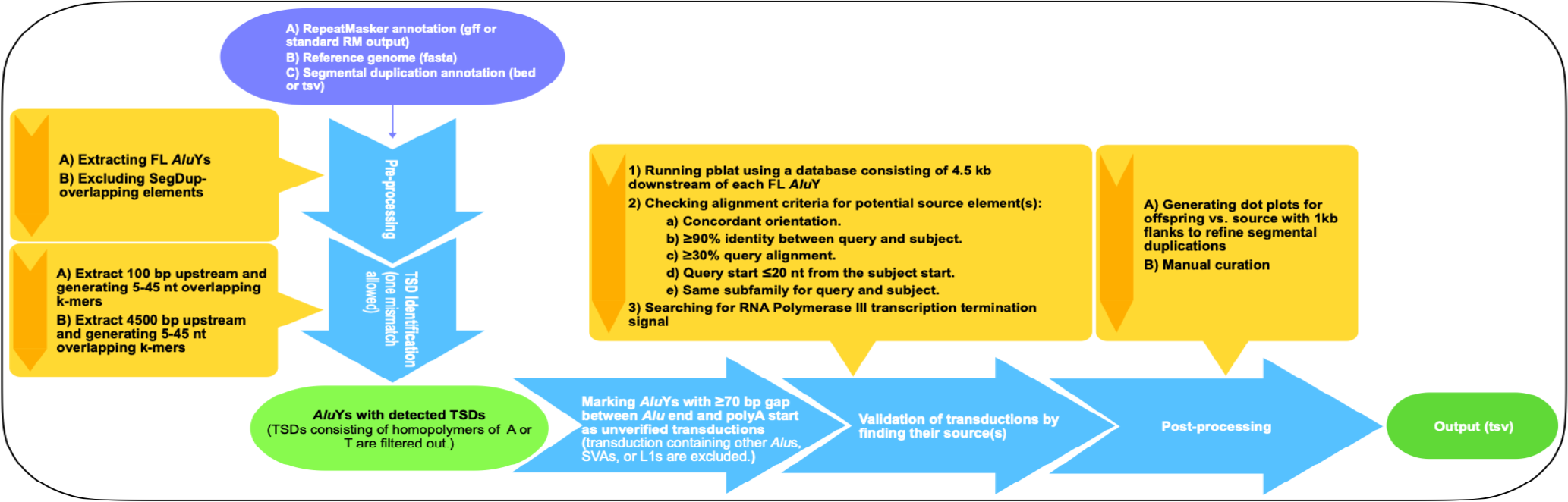
Schematic overview of the pipeline used in this study to identify and verify *Alu*Y-mediated transductions.

Our approach begins by extracting full-length (FL) *Alu*Y subfamily members from RepeatMasker annotations. FL *Alu*Ys are characterized as elements starting within four nucleotides of the consensus sequence and extending to, or beyond, 267 nucleotides (relative to the consensus sequence) in length [3]. The script then filters out FL *Alu*Ys overlapping with segmental duplicates, provided such annotation exists.

For each FL *Alu*Y, our script searches a region spanning 100 bp upstream and 4500 bp downstream to locate TSDs. This involves generating overlapping k-mers (5-45 base pairs in length) and allowing a single mismatch to determine the FL *Alu*Y boundaries. The process includes evaluating the size and position of poly(A) tracts. An *Alu*Y is marked as an unverified potential transduction if the gap between the end of *Alu* and the start of the poly(A) tract is over 70 nucleotides long. Any *Alu*s with a transduction segment comprising another *Alu*, SVA, or L1 element is excluded to avoid any confounding conclusions downstream.

The final step involves transduction validation, applying the following key criteria: (a) aligning each transduced segment to a database of potential source elements using pblat [40] to identify the source element, and (b) ensuring that the transcription termination motif (at least four Ts) is situated downstream of the transduced DNA relative to the source element. This database comprises sequences spanning 4500 bp downstream of each FL *Alu*Y subfamily member. Alignment criteria with pblat include concordant orientation between hit and subject, a minimum 90% identity between query (offspring) and subject (source), alignment of at least 30% of the query (offspring), the alignment start position of the query (offspring) being within 20 nucleotides of the subject (source) start position. Additionally, both offspring (query) and source (subject) must belong to the same subfamily.

To address ambiguities in parent-offspring relationships, particularly in the absence of segmental duplication annotations, we extended our analysis to include 1kb of upstream and downstream sequences of each parent and offspring sequences identified during the previous step. Subsequently, using YASS (command line version) [41], the sequence of each progenitor and its corresponding offspring were aligned against each other, and dot-plots were generated with default settings except for an adjusted E-value of 1e-3. Sequences exhibiting duplicated patterns between offspring and parents were excluded. Ultimately, the final results were subjected to manual curation to generate a *bona fide Alu* transduction catalog.

## 3. Results and Discussion

Many experiments have been conducted to study features of *Alu* element insertions across different genomes. Kojima’s investigation on *Alu* monomers in the human genome revealed eight recent insertions of monomer units, including one originating from another retrotransposed monomer with transduction of a short 3’ flanking sequence [39]. Beyond that, Hoyt et al. identified a new repetitive element termed WaluSat located within the short arms of acrocentric chromosomes of the first complete human genome (T2T-CHM13) [3]. Interestingly, in some cases, these elements were immediately preceded by *Alu*Sx3, both flanked by target site duplications. The authors suggested the possibility of a transduction process contributing to forming these chimeric elements (*Alu*Sx3+WaluSat). Given the absence of prior studies examining genome-wide *Alu*-mediated transductions, in contrast to the extensive analyses of similar events caused by L1s and SVAs, these observations motivated us to investigate the first complete human reference genome (T2T-CHM13) [42, 43] to discover and estimate the frequency of such occurrences mediated by *Alu*s. Specifically, we focused on the most recent *Alu* family, *Alu*Ys, known to harbor active subfamilies capable of ongoing transpositional activity [44] because this ensured high confidence in detecting target site duplications, a prerequisite for the identification of true transductions.

The initial count of *Alu*Y subfamily members from the RepeatMasker output (CHM13v2.0_RM-2022MAR23.out) provided by the T2T consortium was 166,483. However, as described in the Method section, we extracted only 128,695 full-length *Alu*Ys from the RepeatMasker output. Some of these full-length elements are expected to retain their transcriptional and retrotransposition capabilities [3]. In order to ensure the unambiguous assignment of the source element to each offspring, we excluded the full-length *AluY*s that were located within segmental duplication regions [3, 45]. Consequently, our analysis comprised 118,489 full-length *Alu*Ys (Figure 3 and Table 1) as input for our pipeline to detect transduction events mediated by these sequences. While allowing for one mismatch, we found and confirmed TSDs for 118,157 of the analyzed *Alu*Ys (Table 1). The length of TSDs ranged from 5 to 45 nucleotides, with a median of 11 bp (Table 1).

**Table 1.** Comparative summary of *Alu*Y elements and associated target site duplications (TSDs)

| Species (reference genome) | Total count of *Alu*Ys within the RepeatMasker annotation | Full-length *Alu*Ys count | Count of confirmed TSD | Median length of TSDs (bp) |
| --- | --- | --- | --- | --- |
| Human (T2T-CHM13) | 166,483 | 118,489* | 118,157 | 11 |
| Chimpanzee (panTro6) | 131,610 | 106,386 | 105,973 ** | 11 |
| Rhesus monkey (rheMac10) | 234,579 | 191,128 | 190,265 ** | 13 |

\* FL *Alu*Ys located within segmental duplicates are excluded.
\*\* TSDs composed of homopolymers are excluded. Additionally, *Alus* with unknown downstream sequences are filtered out, owing to incomplete genomic sequencing in panTro6 and rheMac10.

**Figure 3.**
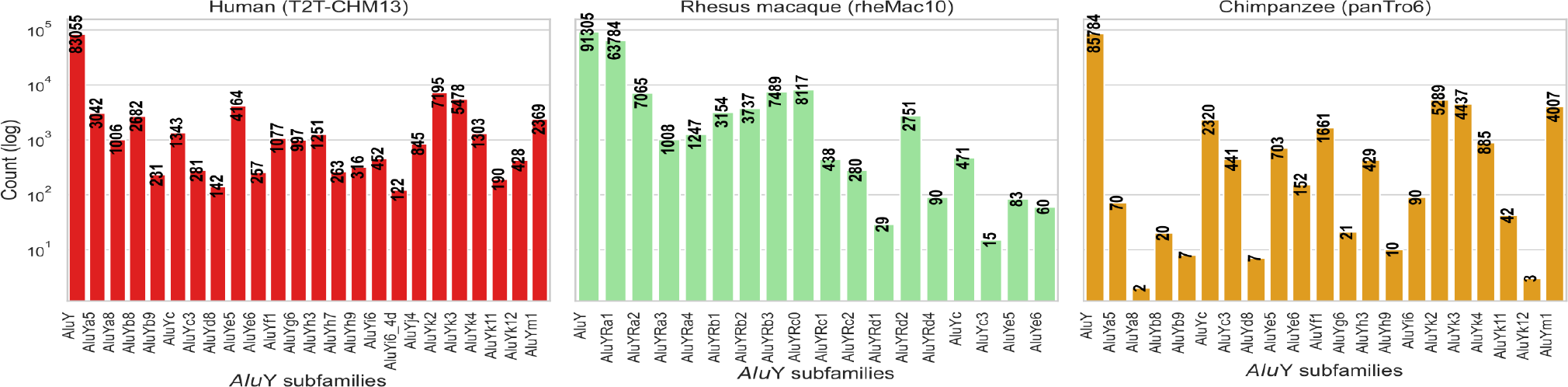
Count of analyzed full-length elements per each *Alu*Y subfamily category.

Of 118,489 FL *Alu*Ys, 633 (0.54%) exhibited a sign of potential DNA transduction. This was indicated by the presence of an additional sequence located between the end of *Alu*, as annotated by RepeatMasker, and the 3’ TSD. Given that our transduction discovery was based on identifying a 3’ TSD located far away from the end of an *Alu*, we used Karlin-Altschul statistics [46, 47], specifically estimating the *E* and *P*-values of their TSDs, in order to ensure these 633 elements represent true transduction events. It is important to note that this step was an extra measure not included in our primary pipeline. The results of this analysis revealed that the estimated values associated with these TSDs were exceptionally high, suggesting that the detected 3’ TSDs, situated distantly from the *Alu* element ends, could potentially be coincidental or random occurrences rather than associated with the transposition mechanism. However, to reduce the likelihood of whether these transduction signatures observed are merely artifacts due to the shortness and low complexity of TSDs (a frequent feature in the genome), we traced the origins of these additional DNA sequences in the 633 *Alu* elements by aligning them to the T2T-CHM13 human reference genome and identifying the presence of an *Alu* transcription termination signal (at least four Ts) after these sequences. Therefore, this validation step was crucial in confirming the initially identified list of transductions and excluding false signals.

While potential sources were identified for 24 of the 633 *Alu*Ys (Table 2), manual examination failed to verify any instances of transductions. Instead, the initially transduced segments identified in the previous step appear to be parts of poly(A) tails that have accumulated mutations, resulting in the formation of microsatellite-like sequences (Table 2). This observation is consistent with prior studies highlighting the contribution of *Alu*s to the generation of satellite-like repeats [15, 48-50]. Moreover, this finding is consistent with studies that suggest slippage by the L1 ORF2 polymerase during insertion can lead to the expansion of the A-rich tail [51, 52], a phenomenon that explains the observed distance of the 3’ TSDs from the end of these 633 *Alu*Ys. The poly(A) tail length variations and sequence heterogeneities suggest the dynamic nature of these regions [49], which add a layer of complexity to the analysis of *Alu* elements.

**Table 2.**
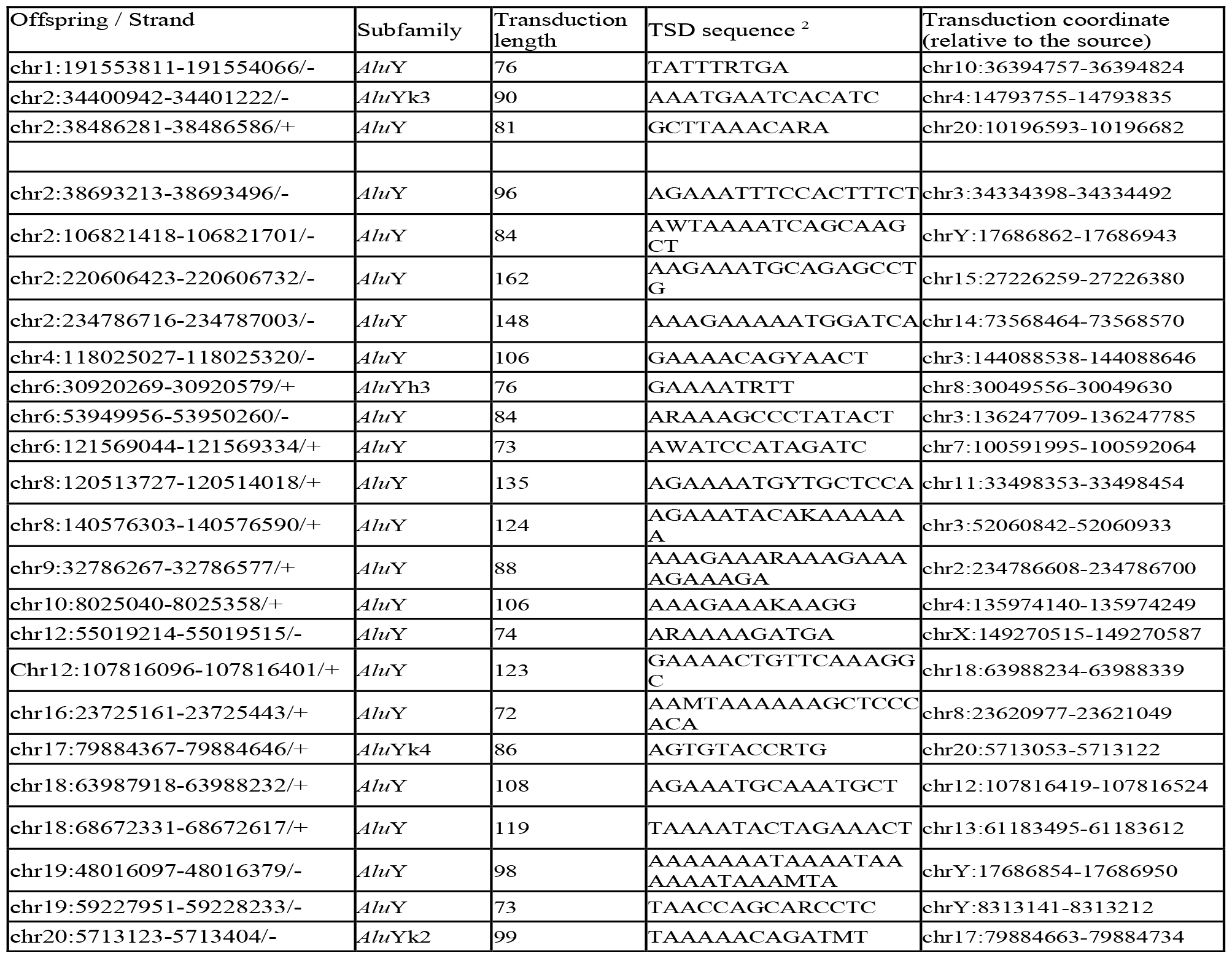
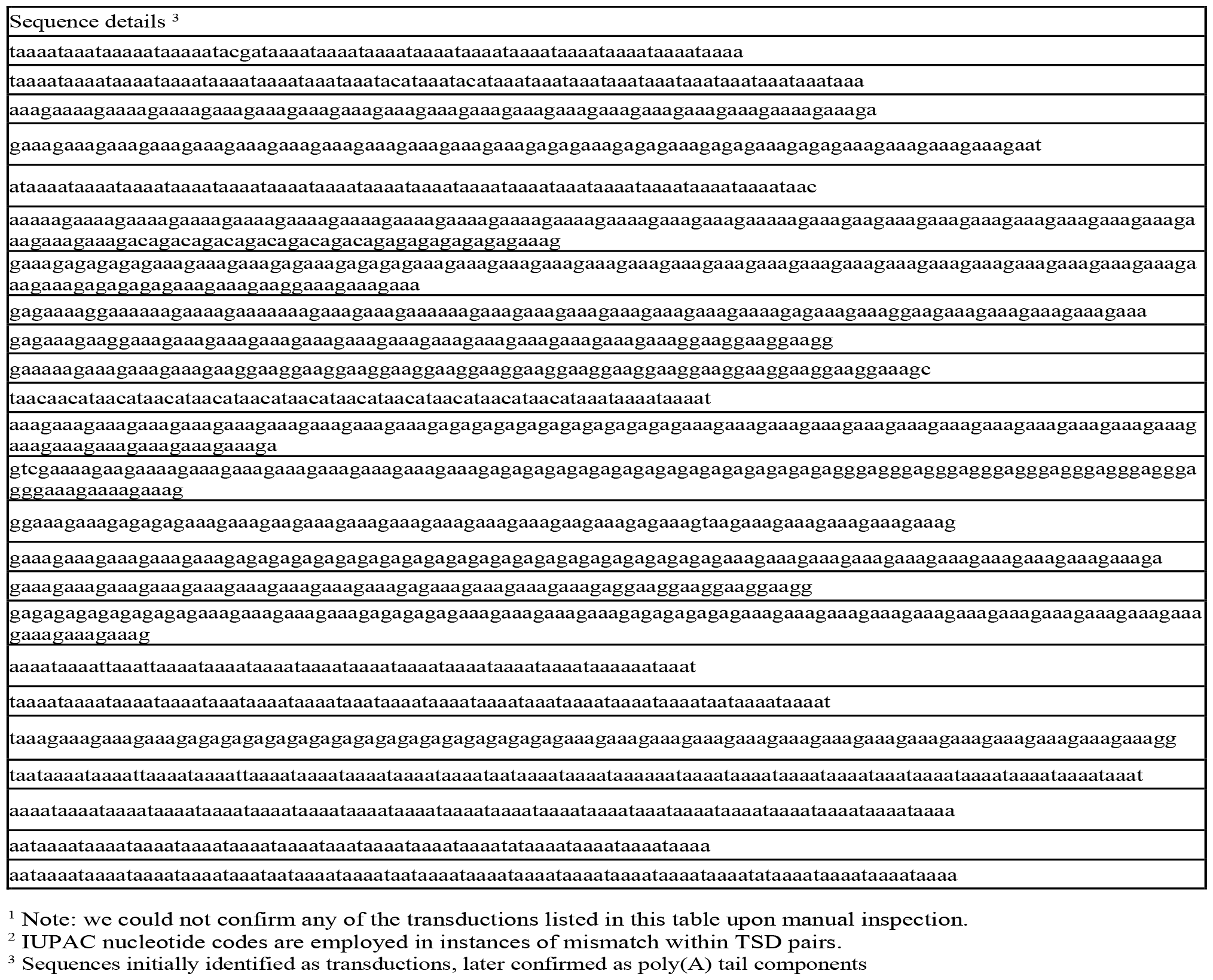
Summary of potential *Alu*Y transductions and source elements within T2T-CHM13 genome^1.^

While our primary focus was the human genome, we extended our scope to non-human primate genomes for a more comprehensive assessment of *Alu*-mediated transductions. Our goal was to investigate whether the infrequency of *Alu*-mediated transductions was unique to humans or a broader phenomenon across other primates in general. Therefore, we analyzed 106,386 *Alu*Y elements in the chimpanzee genome, panTro6, and 191,128 in the rhesus monkey genome, rheMac10, (Table 1 and Figure 1). It is important to highlight that these genomes were not complete telomere-to-telomere assemblies. Consequently, we were unable to examine 162 and 85 FL *Alu*Y elements in the chimpanzee and rhesus monkey genomes, respectively, due to gaps in the sequences downstream of these elements. Initially, approximately 1.3% of the *Alu*Y elements in the panTro6 genome and 1.1% in the rheMac10 genome appeared to display a transduction signature. However, similar to our findings in the human genome, closer inspection revealed that these signatures only consisted of heterogeneous poly(A) tails of varying lengths (Supplementary Tables 1 and 2).

Unlike L1s and SVAs which are transcribed by RNA polymerase II, *Alu*s are predominantly transcribed by RNA polymerase III and thus use a distinct transcription termination mechanism, reflecting their unique interactions with RNA polymerases and subsequent genomic implications. In L1s and SVAs, transcription termination is mediated by a canonical polyadenylation signal (AATAAA) or a variant near the 3’ end of the element, which occasionally is bypassed, resulting in the incorporation of downstream DNA leading to 3’ transduction [3, 34, 45]. In contrast, *Alu* elements employ a termination process characterized by a stretch of at least four thymine bases, oligo(T) [38]. Usually, *Alu* transcripts extend into downstream sequences until they encounter a termination signal, which can result in the inclusion of additional non-*Alu* sequences within their transcripts[15, 53]. However, our study finds that *Alu*-mediated transductions are uncommon in the human genome, unlike transductions rendered by L1 and SVA elements. This finding suggests that the disparity in transduction frequency between *Alu*s and L1s or SVAs can be attributed to differences in post-transcriptional processes, particularly in the integration of the reverse transcript into the genome. It seems that *Alu* transcripts with additional non-*Alu* sequences produced by RNA polymerase III are not optimal substrates for the L1 machinery, impacting their amplification. This is consistent with evidence indicating that *Alu* transcripts with extended non-*Alu* sequences are inefficient templates for retroposition [15, 54]. It is also possible that transcriptionally and retropositionally active *Alu*s (*Alu* master genes) have a strong termination signal 3’ to the element and their transcripts do not contain extra DNA segments. Thus far, master Alu genes have not been identified unequivocally in the human genome, however 88 full-length young *Alu* sequences in the chm13v2.0 reference genome are immediately followed by at least four Ts (Makałowski and Halabian, unpublished data).

In conclusion, our research suggests that *Alu*-mediated transductions in the genomes of human, chimpanzee, and rhesus monkey, are extremely rare. This is a notable deviation from the relatively frequent transductions observed within L1 and SVA elements. It is important to acknowledge that our analysis was centered on the youngest *Alu* elements (*Alu*Ys), which are presumed to be transcribed by RNA polymerase III. Moreover, we applied strict criteria to our analyses to eliminate potential false positives. Thus, the existence of rare *Alu* transductions by co-transcription via RNA polymerase II transcripts or derived from older *Alu* subfamilies, i.e., *Alu*S and *Alu*J elements, cannot be excluded. In summary, despite the fact that *Alu*s outnumber L1s and SVAs, transductions mediated by *Alu* elements seem to be uncommon, if any at all. Moreover, since we analyzed three different primate genomes from a broad phylogenetic distribution, we predict our findings will likely apply to the primate phylum. However, since biology is full of exceptions and surprises, we cannot exclude the fact that rare *Alu*-mediated cases might be discovered in the future.

## Data availability

The codes used for this paper have been deposited on GitHub and can be accessed through https://github.com/IOB-Muenster/

